# Effect of Size Variability, Filamentation and Division Dynamics on Escherichia coli Allometry

**DOI:** 10.1101/2022.12.13.520023

**Authors:** Tanvi Kale, Dhruv Khatri, Chaitanya A. Athale

## Abstract

The cell surface area (SA) increase with volume (V) for cells is determined by growth and regulation of size and shape. Most studies of the rod-shaped model bacterium *Escherichia coli* have focussed on the phenomenology or molecular mechanisms governing such scaling. Here, we proceed to examine the role of population statistics and cell division dynamics in such scaling by a combination of microscopy, image analysis and statistical simulations. We find that while cells sampled from mid-log cultures follow a 2/3 exponent, similar to geometric (Platonic) solids, drug induced filamentous cells have higher exponent values. Modulating the growth rate to change the proportion of filamentous cells, we find SA-V scales with an exponent *>* 2/3, exceeding that predicted by the geometric scaling law. However, since increasing growth rates alter the mean and spread of population cell size distributions, we use statistical modeling to disambiguate between the effect of the mean size and variability. Simulating (i) increasing mean cell length with a constant standard deviation (s.d.) and (ii) a constant mean length with increasing s.d. results in scaling exponents that exceed the 2/3 geometric law, when population variability is included. In order to overcome possible effects of statistical sampling of unsynchronized cell populations, we virtually ‘synchronized’ the estimation of SA-V scaling from single cell growth experiments. We find the exponent depends on the stage with the maximal cell length heterogeneity and scaling exponent observed during the intermediate period between birth (B) and division (D) stages. These results point to a need to consider population statistics and a role for cell growth and division when estimating SA-V scaling of bacterial cells.

## Introduction

Cell shape and size are recognizable characteristics of cellular life forms, and even serve as identification criteria in bacteria. Regulation of bacterial cell shape and size influences how surface area (SA) and volume (V) scale with increasing size. Faster growth results typically in larger cells, as described by Schaechter’s law (Schaechter et al., 1958). On the other hand a small change in shape and SA:V ratio of rod-shaped cells like *Escherichia coli* by filamentation, provides an advantage in terms of increasing the surface area per volume of cell cytoplasm (Koch, 1995; Young, 2006). Thus elongated or filamentous cells can absorb even more nutrients than short rod-shaped cells. This has been observed under nutrient depleted conditions in multiple bacteria, and is thought to be induced by delay or inhibition of septation (Young, 2006). At the same time why a certain cell attains a size is considered an outcome of evolution constrained by the molecular machinery driving it (Bonner, 2006). Since variability in the population is commonly observed due to non-genetic factors, it is important to consider such distributions in any cell shape-size estimations.

Even a population of *E. coli* synchronized, as is now commonly done in microfluidics follow a distribution of sizes with mean values different depending on whether they are in the birth (B) or division (D) stage (Taheri-Araghi et al., 2015). The variability has been variously to variability in either single-cell growth-rates (Hosoda et al., 2011) or cell cycle time (Osella et al., 2016) or stochasticity in DNA replication (Gangan and Athale, 2017). While multiple mechanisms can result in *E. coli* cell filamentation in a population, whether such heterogeneous cell sizes could have a functional role remains unclear.

Single celled organisms take up nutrients through their membranes, and the ratio of the surface area to volume is an important measure of how cell shape and size affect adaptation. For simple geometric objects e.g. spheres or regular cuboids, of length scale *L*, the increase surface area (*L*^2^) with volume (*L*^3^) is expected to scale as *L*^*γ*^ where *γ*, the exponent, is 2/3 or 0.66 – the geometric scaling exponent. Cell shape however does not appear to follow such ideal scaling with a higher exponent observed, explained in part by the diffusion limited nature of nutrient uptake (Koch, 1996). Nutrient uptake in turn is an important selective feature driving specification of cell shape in bacteria (Young, 2006). Indeed the robustness of *E. coli* cell shape is demonstrated by the reversion to typical aspect ratios of length:width in a 50,000 generation long term evolution experiment (LTEE) experiment (Grant et al., 2021). This optimality could be the result of the constraints governing the rate of surface area addition relative to volume growth rate, that sets cell size and aspect ratio, i.e. length:width of a rod-shaped cell (Harris and Theriot, 2016). However, in a recent set of studies it was shown that while gram-negative *E. coli* surface area follows ideal geometric scaling with volume i.e. *γ* ∼ 0.66, *Bacillus subtilis* scales with an exponent of ∼ 0.83 (Ojkic et al., 2019). A growth rate model coupled to FtsZ assembly was developed to explain this scaling based on cell size growth regulation (Ojkic and Banerjee, 2021). However, it remains to be seen if such a difference between two different rod-shaped bacterial species is reproducible given most of the theoretical considerations that suggest greater than geometric scaling.

Here, we have quantified the cell lengths and widths of *E. coli* populations in presence of cell septation inhibitors and different nutrient conditions. The scaling of surface area with volume from the data demonstrates that except for LB, all other conditions result in scaling that are greater than the geometric scaling exponent of 2/3. We examine the role of increasing average cell size and variability for constant averages using statistical simulations and find both processes could explain the SA/V scaling relations measured. The SA/V scaling of single cell growth data to ensure equal representations of all growth stages demonstrates an exponent of ∼0.9. Analysis of cell division cycle stages is used to examine whether variability is cell cycle stage-dependent.

## Results

### Cell surface area scaling with volume and sources of variability

The surface area (SA) of a cell is expected to scale with volume (V) based on the expression:

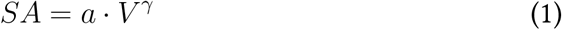

where *γ* is the scaling exponent and *a* is the pre-factor. The size and shape of a cell determines the scaling exponent *γ*. For simple rod-shaped cells a proportionate increase in length and width will result in *γ* ∼ 2/3 ≈ 0.66, as expected when the aspect ratio (*A*_*R*_), i.e. the ratio of length to width, is kept constant (Figure 1A). However, if the cell length increases, while the width of remains constant, for example due intracellular cytoskeletal factors like MreB, or the mechanics of the cell wall, then the scaling exponent is expected to be *γ* = 0.96 (Figure 1A).

**Figure 1:**
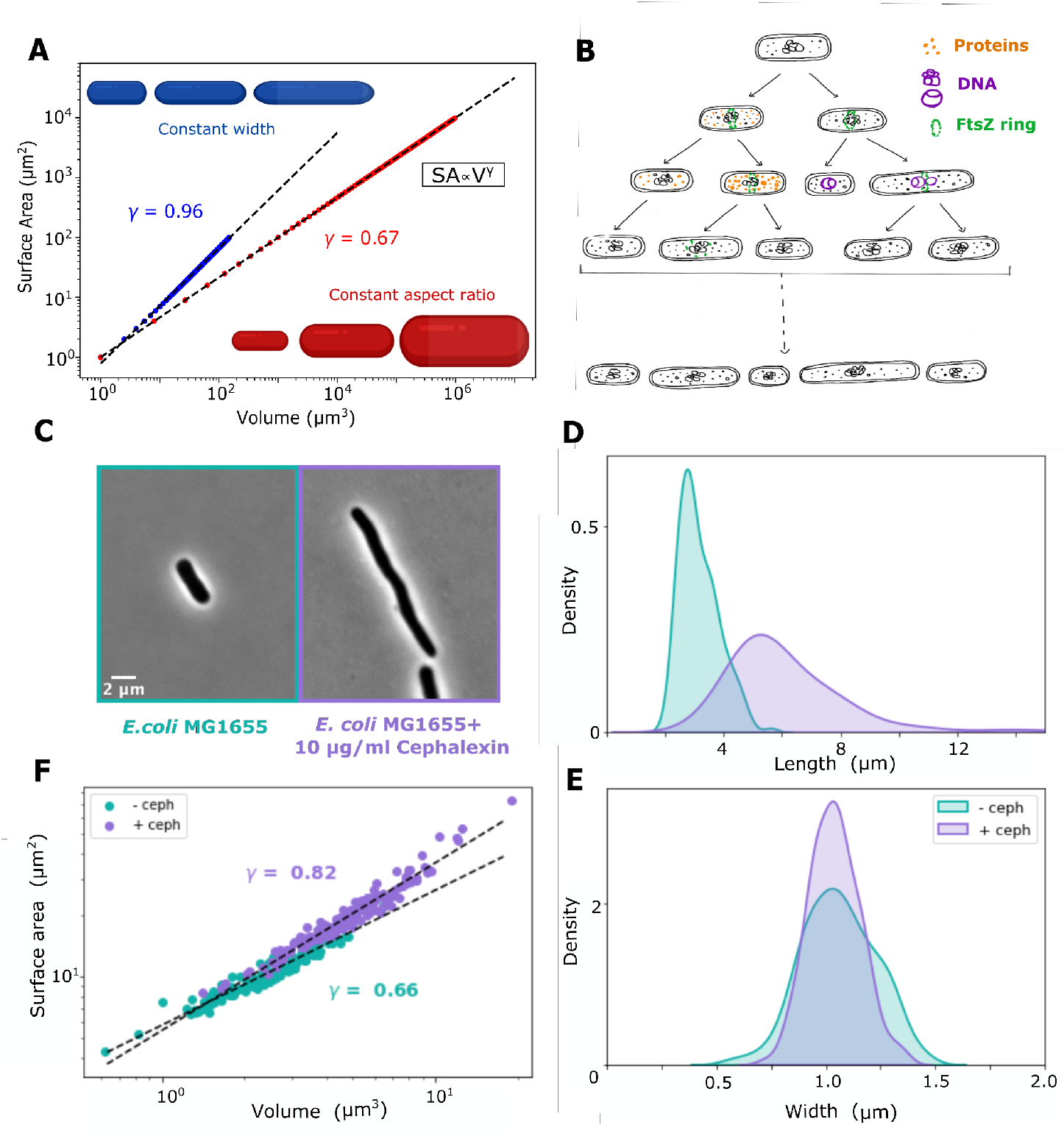
Expected and experimentally observed cell SA-V scaling of *E. coli*. (A) Simulated surface area (SA) and volume (V) data were plotted for rod-shaped bacteria of lengths, L= 1 to 100 *μ*m. Cell Widths (w) are either (i) varied to maintain a constant aspect ratio, *A*_*R*_ = *L/w*=5 (red circles) or (ii) constant *w* = 1 *μ*m (blue circles). Data was fit to Equation 1 (*SA* ∝ *V* ^*γ*^) to estimate *γ*, the scaling exponent. (B) The schematic depicts multiple mechanisms that lead differences in cell size: asymmetric protein segregation (orange), DNA replication stochasticity (purple) and inhibition of the FtsZ ring (green), during multiple rounds of bacterial cell division. (C-F) Microscopy and quantification of *E. coli* cell sizes. Representative images of untreated and 10 *μ*g/ml cephalexin treated *E. coli* are seen in phase contrast microscopy. (D) Cell length and (E) width density distributions are plotted from untreated (n=160) and cephalexin treated (n=111) conditions. (F) The surface area (*SA*) is plotted as a function of volume (*V*) for each cell analyzed (see Materials and Methods) on a log-log plot and fit to Equation 1. The scaling exponent *γ* is the fit parameter. Each circle denotes a single cell. Colors indicate untreated (cyan) and cephalexin treated cells (purple). n: number of cells analyzed.

In a population, cell size variability in terms of different lengths, and the presence of filamentous (i.e. elongated) cells arises from multiple factors or their combination such as single cell gene expression stochasticity (Elowitz et al., 2002), variability of molecular partitioning at division (Rosenfeld et al., 2005), replication stochasticity and coupling to division (Gangan and Athale, 2017) or DNA damage response (Raghunathan et al., 2020) as summarized in Figure 1B. At the same time the variability of widths has been variously reported to be constant (Taheri-Araghi et al., 2015).

It remains unclear whether this variation in widths and lengths could affects the estimation of the scaling exponent. In order to examine this, we first examined SA-V scaling in filamentous *E. coli*.

### Scaling of SA-V of populations of filamentous *E. coli* deviates from the 2/3 law

*E. coli* MG1655 cells were imaged in phase contrast (label-free) microscopy and as expected cells treated cephalexin (10 *μ*g/ml) were filamentous and compared to them untreated cells were shorter (Figure 1C). An image analysis pipeline was used to automatically estimate cell lengths and widths from such label-free images (Materials and Methods). Cell lengths of *E. coli* differ between treated and untreated both in terms of mean and spread of the distribution (Figure 1D). Cell widths on the other hand of both strains have the same mean values around which the data is symmetrically distributed (Figure 1E). The surface area, SA (Equation 2) and volume, V (Equation 3) of individual cells were calculated from the lengths and widths (Materials and Methods). The plot of SA with V was fit to Equation 1 to estimate *γ*, the scaling exponent (Figure 1F). The scaling exponent of untreated cells is *γ* = 0.66, while filamentous cells have *γ* = 0.82.

We infer that filamentation results in a deviation from the ideal geometric scaling of *E. coli* SA with V. In order to test whether intermediate distributions of cell lengths also show this effect, we modulated nutrient availability since it has been shown to change the cell length distribution of populations.

### Increasing growth rate results in population scaling exponents intermediate to filamentous cells

We tested the effect of modulating growth rates on cell sizes by growing them in minimal medium (M9 salts) supplemented with carbon sources- either 4% sorbitol, glycerol or glucose (Materials and Methods). While qualitatively the cells appear similar in size (Figure 2A), the population distribution of cell lengths demonstrates highest modal lengths in M9+Glucose, while the tail of the distribution in M9+Glycerol was longer than M9+Sorbitol, but their modes were comparable (Figure 2B). Cell width distributions in all 3 conditions were comparable (Figure 2C). The SA scales with V in all three media with exponents of 0.7 to 0.8 (Figure 2D). The exponent is apparently linearly correlated to the bulk growth rate (Figure 2E).

**Figure 2:**
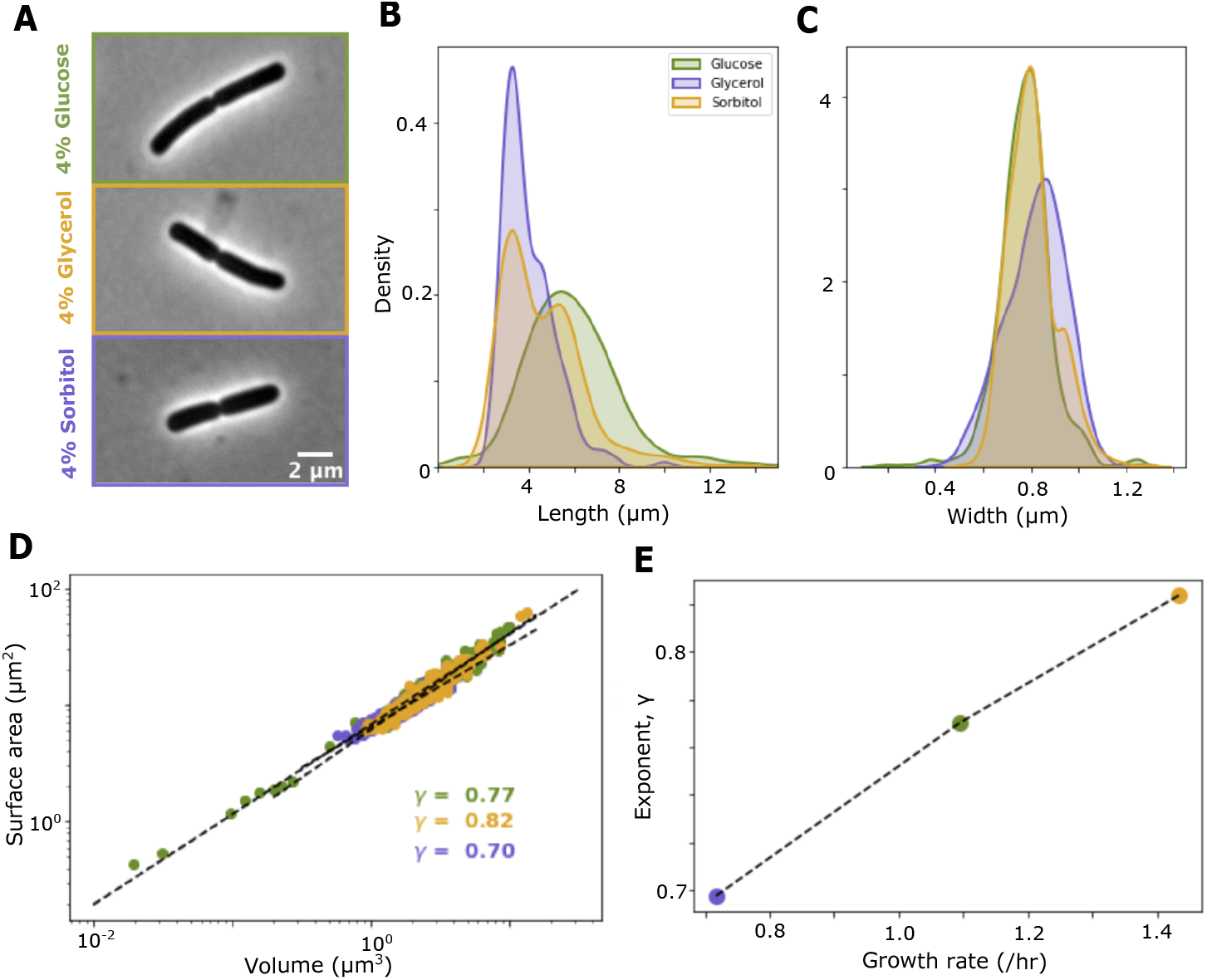
SA-V Scaling exponents of *E. coli* populations with increasing growth rates. (A) Representative phase contrast images of *E. coli* MG1655 grown in M9 supplemented with either glucose (n = 412), glycerol (n = 348) or sorbitol (n = 449) (4% each) were quantified to obtain (B) cell length and (C) width distributions. The surface area and volume calculated from the lengths and widths (circles: individual cells) were fit to Equation 1 obtain scaling exponents *γ* for each growth condition. (E) The *γ* values obtained for the different growth media are plotted as a function of respective growth rates. Colors indicate carbon-source: glucose (green), sorbitol (yellow) and glycerol (purple) used to supplement the M9-medium (as described in the Materials and Methods). n indicates the number of cell analyzed.

Cell length differences from modulating the growth rate are seen both in terms of the mean and the spread of the distribution. In order to distinguish which of these influences the change in the SA-V scaling exponent, we used statistical simulations, since we were unable to separate the two effects experimentally.

### Simulating the effect of increasing mean cell size and variability on SA-V scaling

Controlling cell length mean and standard deviations independently in experiments is technically challenging, therefore we use statistical simulations to address whether variability or means of lengths affect the scaling exponent. To this end we modeled two scenarios:

*Model 1: Increasing mean length with constant variability (sd):* Motivated by Schaechter’s law and our own experiments, where increasing growth rate results in increased mean cell lengths, we examine 1 to 12 *μ*m. In order to test the effect of increasing the mean and a proportionate standard deviation, We simulated cell lengths by sampling a lognormal distribution with increasing means from 0.1 to 2.5 *μ*m and constant standard deviations (Figure 3A). The widths were simulated by sampling from normal distributions with their means increasing proportionately, based on a constant aspect ratio, *A*_*r*_ of 5 (Figure 3B). These length and width distributions result in mean and s.d. values comparable in range to experimental data (Table 2). We find the SA-V of such simulated data scales with correspondingly increasing *γ* values ranging from 0.65 for for short cells where *μ*_*L*_ = 0.1 *μ*m to 0.93 for long cells where *μ*_*L*_ = 2.5 *μ*m (Figure 3C). Averaging of SA and V, that eliminates population variability, results in the recovery of the geometric scaling exponent of *γ* = 0.67 (Figure 3C inset). This demonstrates that even though the mean length and width increased proportionately (constant aspect ratio), the population variability governs the scaling relation. Thus averaging, that eliminates the variability, and the SA-V scaling exponent reverts to 2/3. When we maintain the distribution data, the increasing mean also corresponds to an increase in the exponent (Figure 3D).

**Figure 3:**
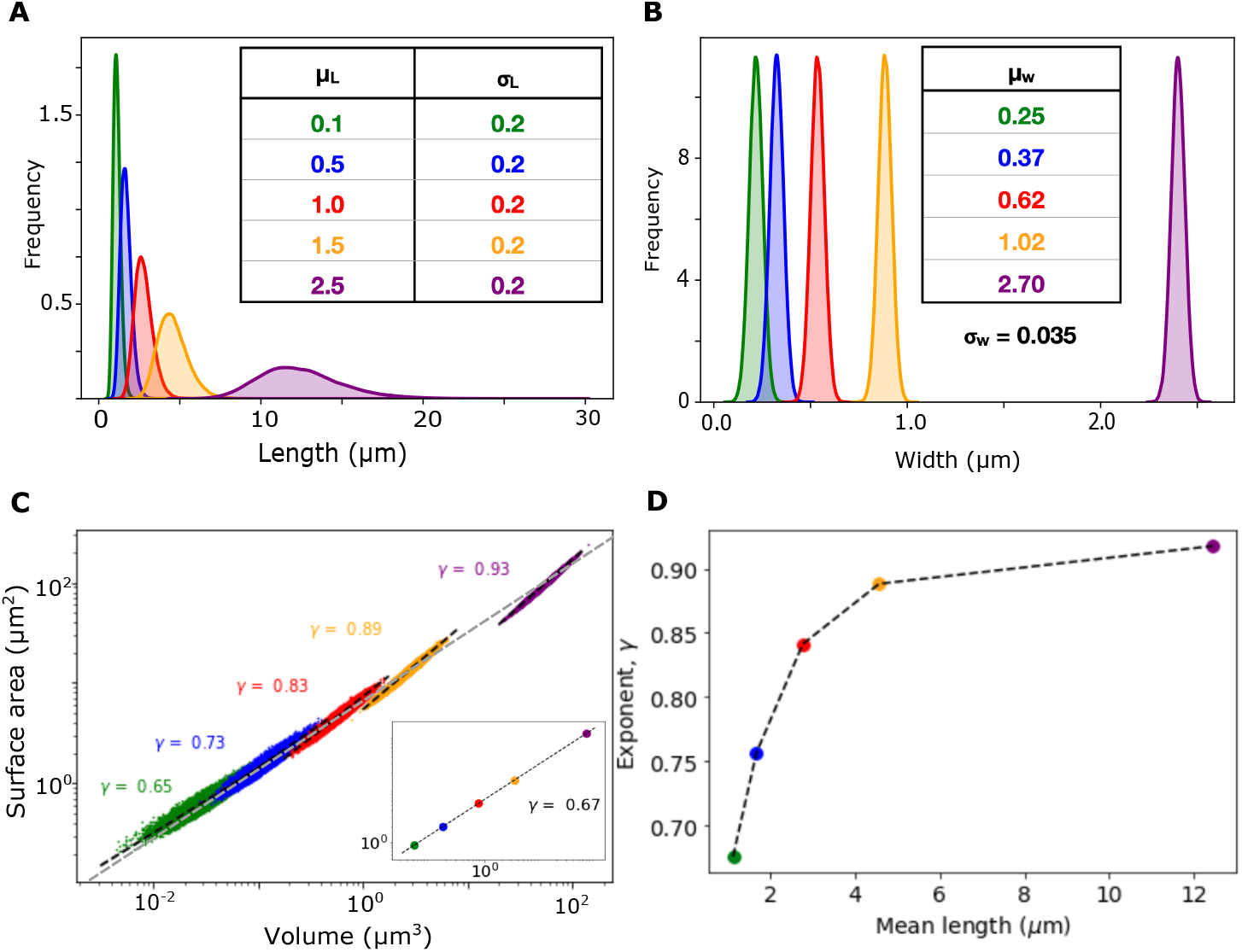
Simulating the effect of increasing mean cell size on SA-V scaling. (A,B) Cell length and width distributions were simulated based on (A) lognormal distributions for a range of mean lengths (*μ*_*L*_) with s.d. (*σ*_*L*_) and (B) a normal distribution of widths. The simulated widths have a mean (*μ*_*w*_) that is determined by *μ*_*w*_ = *μ*_*L*_*/A*_*R*_, where *A*_*R*_ is the aspect ratio (L/w), taken to be *A*_*R*_ = 5, based on experiments. The width s.d. (*σ*_*w*_) is 0.035. (C) The surface area and volumes calculated for the each simulated distribution pair of lengths-widths were fit to to the scaling law (Equation 1, Materials and Methods) to obtain multiple values of the scaling exponent scaling exponent *γ*. Circles indicate one simulated cell. (inset) This is compared to to the single exponent fit to average SA and V from each distribution. Each colored circle is the average of the distribution. (D) The arithmetic mean length of each distribution plotted as a function of the exponent obtained for the distribution. Colors indicate the simulated dataset of cell lengths and the corresponding widths. Number of simulated points per distribution are n=10^6^

*Model 2: Increasing variability (sd) for constant mean length:* In order to address whether the increase in *γ* seen in the earlier simulations can be attributed to spread alone, we simulated distributions with identical mean cell lengths (*μ*_*L*_ = 1 *μ*m) and increasing standard deviation from 0.1 to 0.5 *μ*m (Figure 4A). Widths were sampled as before from a normal distribution with a constant mean (Figure 4B), based on a constant (mean) aspect ratio of 5. We find the SA scales with V with increasing exponents ranging from *γ* of 0.72 to 0.89 (Figure 4C). Once more averaging the SA and V data from the population to get just one average value of SA and V for each distribution, resulted in the geometric scaling exponent of *γ* = 0.67. Interestingly we find the increase in the exponent with CV of length (CV_*L*_) is qualitatively similar to that observed in exeperiment (Figure 4D), but not a quantitative match. Simulated distributions result in a saturation of *γ* with increasing CV_*L*_ (Figure 4E).

**Figure 4:**
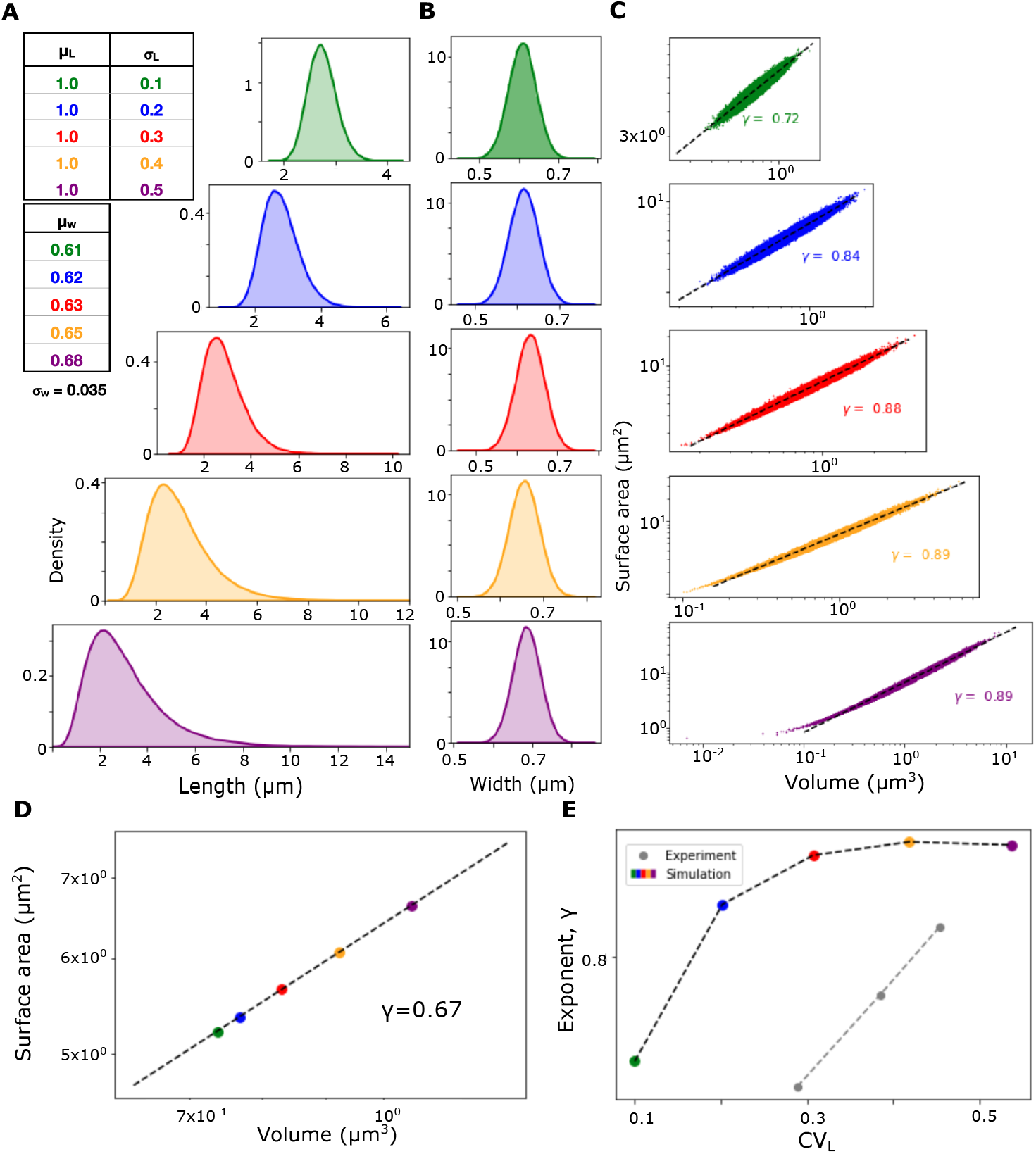
Simulating the effect of constant mean cell length and increasing variability on SA-V scaling. (A,B) Cell size variability was simulated such that (A) cell-lengths follow a lognormal distribution with a constant mean and increasing s.d., (B) while widths follow a normal distribution. (*Boxes*) Mean length (*μ*_*L*_) and s.d. (*σ*_*L*_) and mean widths (*μ*_*w*_) used as inputs to the simulation. (C) SA plotted as a function of V with each point representing a simulated cell sampled from the distributions, was fit to the scaling relation (Equation 1) to obtain the exponent *γ*. (D) A plot of average SA and V obtained for each distribution of lengths-widths is fit to Equation 1 to estimate *γ*. (E) The *γ* values from each of the distributions are plotted as a function of the coefficient of variation of the length (based on the output mean lengths and s.d. as seen in Table 2). Colors: cell length distributions, with their corresponding width distributions. n = 10^6^ for each distribution.

Based on the results from experiment and simulation, variability appears to play an important role in estimating the exponent of SA-V scaling. Since experimental data was sampled from an unsynchronized population, we proceeded to test whether the statistics would still hold true for single cell data.

### Single cell growth dynamics result in SA-V scaling by an exponent ∼0.9

We grew *E. coli* MG1655 expressing GFP on nutrient agar pads and imaged them in fluorescence (Materials and Methods). Cells were visualized by expressing the fluorescent protein GFP from a plasmid. We used a MATLAB based image-analysis pipeline to track representative cells from image time series (Figure 5A, Supplementary Video SV1A-C) to obtain length and width dynamics. The plot of SA scales with V for each individual cell with the progress of the cell division cycle with an exponent of *γ* between 0.8 and 0.9 (Figure 5B). We observe a similar value for *γ* on fitting SA vs V to Equation 1 for all the cells together (5C). This appears to suggest that the SA of individual cells also scales with V with an exponent exceeding the geometric 2/3 exponent, consistent with population sampled data and simulations, with a mean *γ* value of 8.2 (5D).

**Figure 5:**
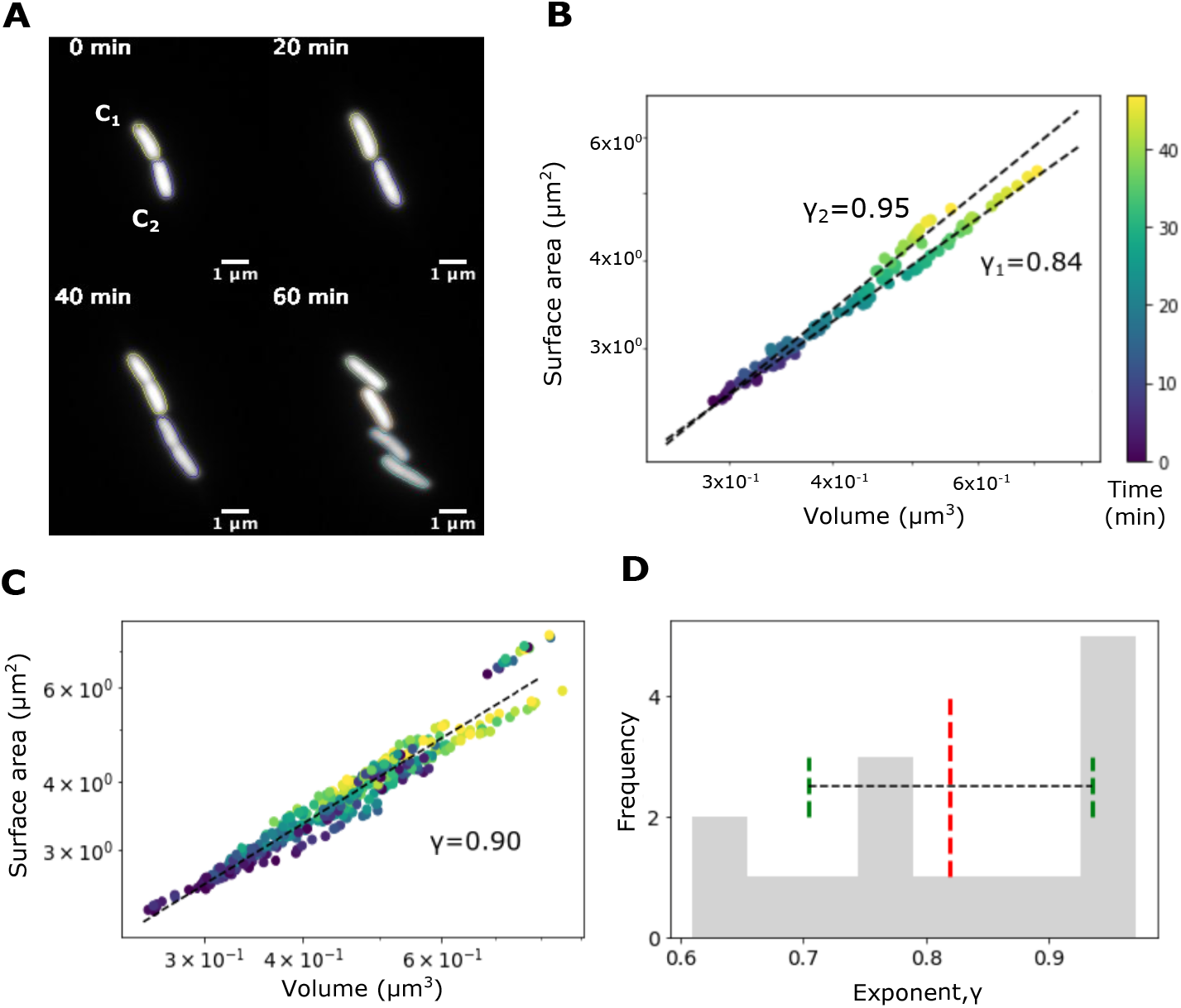
Effect of single cell growth dynamics on SA scaling with V. (A) Images of *E. coli* MG1655 expressing GFP from a plasmid grown on agarose pads show individual time-points from time-lapse microscopy. Time: min. Scale-bar: 1 *μ*m. (B, C) The SA was plotted as a function of V with time encoded in color (B) for two representative daughter cells (C1 and C2) with scaling exponents *γ* of 0.83 and 0.94 respectively and (C) for 16 single-cell time series fit to one line resulting in *γ*=0.9. (D) Frequency histogram depicts the distribution of exponents obtained when each time-series was fit individually. Mean (red dashed line) and s.d. (green dashed lines) of the exponents are also represented.

### Effect of cell length variability dynamics during *E. coli* cell growth on SA-V scaling

The time from birth to division of single cells was used as a measure of single-cell life span (Figure 6A). We divided the normalized lifespan (scaled from 0 to 1) into four equal segments that we refer to as birth (B), C1, C2 and division (D) stages (Figure 6B). As expected, the mean of the cell length increases with ‘stage’ of cell division (Figure 6C), while the width appears to have a constant mean but slight increase in spread (Figure 6D). This data is used to calculate the SA-V scaling of individual cells through the cell cycle stages resulting in exponents that change within the lifetime of cells (Figure 6E). Surprisingly, we find the change exponent *γ* and variability of lenths (CV_*L*_) are highest in stages C1, C2 with *γ* ∼0.8) and at division *γ* ∼0.66, the geometric exponent (Figure 6F). This suggests that cell size variability changes during the process of growth and division and is consistent with population statistics.

**Figure 6:**
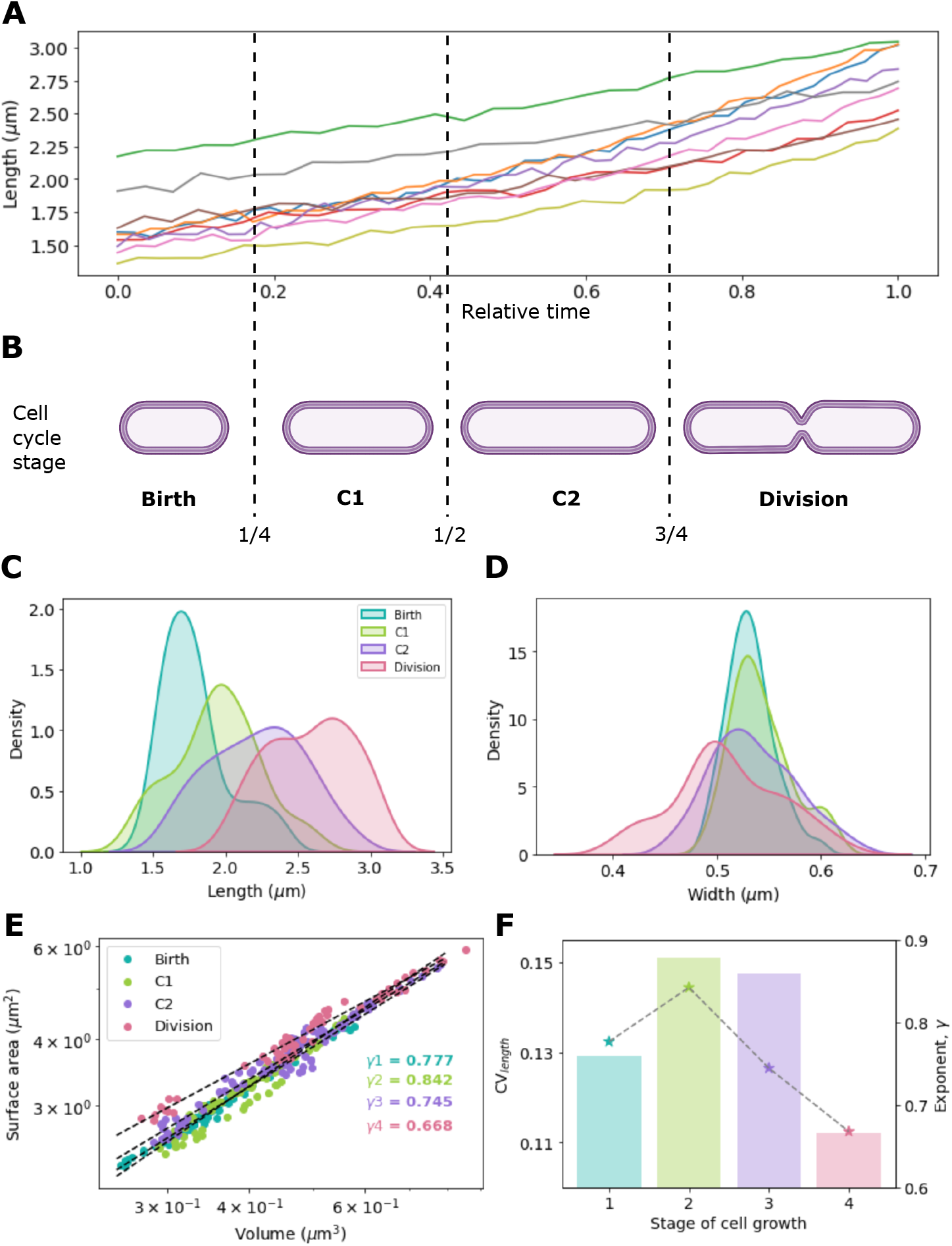
Effect of growth dynamics of single cells on variability and SA/V scaling. (A) The length (*μ*m) as a function of relative time in the cell cycle is used to identify (B) four ‘stages’ of cell growth: birth, c1, c2 and division. (C, D) The stage dependent frequency distribution of (C) cell lengths and (D) widths are plotted. SA is plotted as a function of V for individual cells. Each point represents an individual cell (n = 10). Colors represent the stage of the cell division cycle. The data was fit to the scaling relation to infer *γ*. (F) The cell cycle stage dependent coefficient of variation of cell length, CV_*L*_, (bars) and SA/V scaling exponent *γ* (*) and are plotted.

## Discussion

The scaling of surface area and volume of cells is fundamental to comparative physiology and could offer insights into regulatory mechanisms at cellular scales. In this study we find that in populations of *E. coli* cell surface area scales with volume with an exponent of *γ* ∼ 0.6 when cells are rapidly growing and sampled from mid-log phase. However, drug-induced filamentation or a range of sugars that modulate growth rate, result in higher exponents between 0.7 to 0.8, i.e. consistently higher than 2/3. Statistical simulations where either the mean cell length or the variability in cell length is systematically increased for distributions of cell lengths and widths result in SA-V scaling with an exponent greater than 2/3. Averaging the simulated SA and V distributions results in a return to the 2/3 scaling in simulations, demonstrating that population cell variability can alter the inference of the allometric scaling exponent of cell surface area with volume. In order to test whether the lack of synchrony in populations had influenced our scaling exponent estimation, we measured SA-V scaling in growing single-cells and found *γ* ∼ 0.7 to 0.9 during the life-span of a single cell, suggesting such scaling is intrinsic to *E. coli* growth and is independent of sampling. Finally we divided the cell division cycle into equal 1/4th parts and find cell length heterogeneity and by extension the geometric scaling exponent changes from ∼0.6 to ∼0.8, depending on the cell division stage. We find inter-division stage SA-V scaling to have the highest exponent, while dividing cells follow the geometric law (2/3), suggesting relevance for cell size control mechanisms.

In our experimental analysis, we have shown that the measured scaling exponent of SA-V scales linearly with the growth rate (Figure 2E). In contrast simulated scaling exponents saturate with cell length variability (Figure 4E). This could be explained by the fact that in simulations the cell shape as measured by the mean aspect ratio (length:width) is held constant. However, it appears that in experiment, cell length increase does not correspond to a proportionate change in cell width. This suggests that in order to obtain a better match of SA-V allometric relations between our model and experiments, we will need an improved model of cell shape regulation where a mechanical feedback between membrane curvature and envelope expansion maintain cell shape (Al-Mosleh et al., 2022). An addition to such a model would be the reconciliation of population data of the kind presented here.

The functional and biological relevance of SA-V scaling, has been discussed previously in terms of its effect on the surface area to volume ratio (*SA/V*), with suggestions for higher than the 2/3 law based on benefit to uptake of resources (Huxley et al., 1993; Okie, 2013). We consider this for the rod-shaped cell geometry of cell populations with increasing sizes or higher variability, e.g. presence of filamentous cells. The area per unit volume (SA/V ratio) decreases with increasing size, based on geometry. Since nutrient uptake is required to cross the surface-barrier of a cell, the uptake rate per unit area, or flux, can be written as being proportional to area as *Flux ∝ SA*. By taking the product with 1*/V* on both sides we arrive at 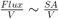. Assuming V to be independently increasing, corresponding to larger cells, an increase in SA will be based on Equation 2 and the SA-V scaling exponent *γ*. A *γ* value of 2/3 (or 0.66) will scale slower than 0.7 or 0.8, suggesting that flux will be higher for cells of the same length scale whose SA-V scale with *γ >* 2/3. This could suggest a potential functional role for the observed exponents in *E. coli* cells.

Deviation of unicellular bacteria from the geometric 2/3 law is expected due to ecology and survival (Okie, 2013). Indeed predictions of phenomenological models suggest cells may optimize their shape and size within the constraints governing the rate of surface area addition relative to volume growth rate (Harris and Theriot, 2016). However, a recent set of studies proposed gram-negative *E. coli* surface area follows ideal geometric scaling with volume, *γ* ∼ 2/3, while for *B. subtilis* it is ∼ 0.83, that was interpreted as being indicative of alternative mechanisms at play in cell size growth regulation (Ojkic et al., 2019). A growth rate model coupled to FtsZ assembly was developed to explain this difference in scaling (Ojkic and Banerjee, 2021). In our data, we too find rapidly growing cells, potentially sampled in division phase, do scale with an exponent of 2/3. However, we find the when either other growth conditions are used for sampling population cell sizes, or dynamics of single cell growth are examined, the exponent is consistently higher than the geometric law. Our approach demonstrates could be used to provide further inputs to disambiguate between multiple phenomenological models of bacterial cell size regulation such as ‘sizer’, ‘timer’ and ‘adder’.

In conclusion, we find SA-V scaling exponents of single cells of *E. coli* to exceed the geometric exponent value of 2/3 in a manner that depends on drug-induced filamentation, varying growth rate and cell growth dynamics. Statistical simulations demonstrate that without the inclusion of population variability the SA-V scaling exponent reverts to the geometric law. We find, that the dynamics of single cell growth in multiple experiments, recapitulates the scaling exponent measured in population ensembles and provides quantitative data that can be compared to phenomenological and mechanistic models of bacterial cell division.

## Materials and Methods

### Bacterial culture and sample preparation

Analysis of populations of cells sampled from mid-log involved growing *E. coli* MG1655 in multiple liquid media to modulate their growth rates- Lysogeny Broth (LB), M9-salts supplemented with glucose (4% w/v), sorbitol (4% w/v) and glycerol (4% v/v) (Fisher Scientific, Mumbai). Growth rates were measured by monitoring optical density at 600 nm (OD600) every 10 min in a multiwell plate reader (Varioskan Flash2, Thermo Fisher Scientific, Waltham, MA, USA) and fitting the data to a logistic function, as described previously (Gangan and Athale, 2017). At mid log-phase, cells were sampled, washed three times with 1% sterile phosphate-buffered saline (PBS) and mounted on slides coated with 10 *μ*l of 1% (w/v) low-melt agarose (HiMedia, Mumbai, India) and observed in microscopy.

Single-cell growth dynamics was performed using *E. coli* MG1655 transformed using the Hanahan’s method of CaCl_2_ chemical competence and heat shock (Sambrook and Russell, 2001) with a GFP expression plasmid based on a pET15b backbone with a P_*lac*−*ara*_ hybrid promoter (AND-gate). Cells were grown overnight in LB Broth at 37°*C*, of which 10 *μ*l spread on an 2×2 *cm*^2^ agarose pad made on a slide using 1.5% Agarose in LB Broth supplemented with 100 *μ*M IPTG and 0.7% (w/v) arabinose. A coverslip with the agarose pad was placed on a custom-built 3D printed holder for time-lapse microscopy.

### Microscopy

#### Single time point microscopy of populations

Cells grown in growth media were washed in phosphate-buffered saline (PBS) and a 5 *μ*l droplet was placed on an agarose-pad and imaged in 100x (N.A. 1.45) phase contrast mode using an Olympus upright microscope with Olympus cellSense Dimension 3.1 software for image acquisition (Olympus Corp., Japan) and a CCD camera Hamamatsu ORCAflash4.0 (Hamamatsu Photonics, Japan).

#### Time-lapse microscopy of single cells

Cells were mounted between two coverslips and imaged using a 100x N.A. 1.4 objective in fluorescence (excitation/emission 488/510 nm) for GFP mounted on a Nikon TiE inverted epifluorescence microscope illuminated with a 130W Intensilight Optic Illumination with control software NIS Elements ver.4.30.01 (Nikon Corp., Japan). Images were acquired on a CCD camera Andor Clara 2 (Andor, NI, U.K). Images were acquired at 1 minute intervals for 60 minutes.

#### Image analysis

All low level image intensity adjustment, selection of regions of interest (ROI) and adding scale-bars was performed using Fiji (Schindelin et al., 2012).

In order to detect cell outlines, we developed an analysis pipeline in MATLAB (Mathworks Inc., Natick, MA, USA) using functions provided through the Image Processing Toolbox. The workflow was as follows: (a) Cell outlines were identified by a combination of edge-detection by the Canny method (Canny, 1986), gaps filling, and regions bridging. (b) Cells were filtered based on a minimum solidity criterion between 0.3 and 1 to remove small and irregular objects, with cell boundaries were optimized using an edge-based active contour method (Kass et al., 1988), with a smoothing factor 1 and a contraction bias of -0.1. (c) Optimized contour boundaries were overlaid on the original input image and artefacts were interactively deselected to avoid cell debris and out-of-focus cells. (d) The remaining cells with contours were analyzed for length and width by first detecting the midline, and widths taken from averaging the 3rd and 4th quartiles of the minimal distance between the midlines and the contour boundaries. By taking only the top 50% values we exclude narrow widths at the ends of the cell from the analysis.

The code is available as OpenSource and can be downloaded from https://github.com/CyCelsLab/bacterialsizeScaling. The mean, standard deviation and the CV of the obtained lengths and widths is reported in Table 1

**Table 1:** Experimentally measured statistics of *E. coli* MG1655 cell sizes under multiple growth conditions. The mean ± standard deviation (s.d.) and median values cell lengths and widths measured in experiment are reported here. These correspond to the data plotted in Figures 1 and 2. LB: Lysogeny Broth and M9: minimal medium.

| Growth condition/Media | Mean ± s.d. | Median | CV ($\sigma/\mu$) |
| --- | --- | --- | --- |
| *Length* |  |  |  |
| LB | $3.15 \pm 0.67$ | 2.90 | 0.21 |
| LB + 10 $\mu\text{g/ml}$ cephalixin | $6.34 \pm 2.61$ | 5.05 | 0.41 |
| M9 + 0.4% Glucose | $6.00 \pm 2.31$ | 5.73 | 0.38 |
| M9 + 0.4% Glycerol | $4.00 \pm 1.15$ | 3.63 | 0.28 |
| M9 + 0.4% Sorbitol | $4.71 \pm 2.14$ | 4.22 | 0.45 |
| *Width* |  |  |  |
| LB | $1.05 \pm 0.16$ | 1.04 | 0.15 |
| LB + 10 $\mu\text{g/ml}$ cephalixin | $1.04 \pm 0.11$ | 1.03 | 0.10 |
| M9 + 0.4% Glucose | $0.77 \pm 0.11$ | 0.77 | 0.14 |
| M9 + 0.4% Glycerol | $0.81 \pm 0.12$ | 0.83 | 0.14 |
| M9 + 0.4% Sorbitol | $0.80 \pm 0.10$ | 0.79 | 0.12 |

**Table 2:** Parameters used in the statistical simulations. Simulations of cell size lengths used INPUT lognormal mean (LM) length (*μ*_*L*_) and standard deviations (*σ*_*L*_) resulting in OUTPUT values that are arithmetic mean (AM) and s.d. The values correspond to data plotted in Figures 3 and 4. The range of values used in the simulations correspond to the two models tested: (i) increasing values of *μ*_*L*_ for constant *σ*_*L*_ (ii) a range of *σ*_*L*_ for a constant *μ*_*L*_. The models are described in detail in the Materials and Methods section,

| INPUT |  | OUTPUT |  |
| --- | --- | --- | --- |
| $\mu_L$ | $\sigma_L$ | $\mu_L$ | $\sigma_L$ |
| *(i) Model of increasing means, constant s.d.* |  |  |  |
| 0.1 | 0.2 | 1.13 | 0.23 |
| 0.5 | 0.2 | 1.68 | 0.34 |
| 1 | 0.2 | 2.77 | 0.56 |
| 1.5 | 0.2 | 4.57 | 0.93 |
| 2.5 | 0.2 | 12.44 | 2.53 |
| *(ii) Model of constant mean, increasing s.d.* |  |  |  |
| 1 | 0.1 | 2.73 | 0.27 |
| 1 | 0.2 | 2.77 | 0.56 |
| 1 | 0.3 | 2.84 | 0.87 |
| 1 | 0.4 | 2.94 | 1.22 |
| 1 | 0.5 | 3.08 | 1.64 |

This approach was used for both single time-point images in phase contrast as well as fluorescence images of *E. coli* expressing GFP with small changes in parameters for detection.

### Data analysis

Bacterial growth rates in different media were estimated by fitting mean data obtained from triplicate measurements of optical density at 600 nm every 10 minutes for 3 hours to a logistic model. Comparable to previous work, we assume the cell geometry to be a spherocylinder. This allows us to use the cell length and width to calculate the surface area (SA) as:

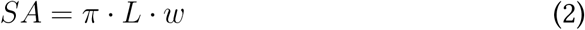

while the volume (V) is given by:

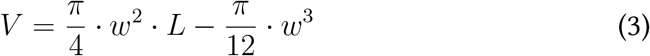

*L* is the pole-to-pole length and *w* the cell width. The scaling relation between surface area (SA) and volume (V) was obtained by fitting the data to the scaling relation in Equation 1. The aspect ratio (*A*_*R*_) was calculated as *A*_*R*_ = *L/w* for each data point. In-house developed Python scripts were used to perform the calculations (Python3) using Pandas, NumPy, SciPy packages and Matplotlib (v3.4.2), with *polyflt* used for curve-fitting (Harris et al., 2020).

The coefficient of variation (CV) was calculated as standard deviation/mean for the simulated distributions and length distributions from experiments.

## Acknowledgments

We are grateful to Nishad Matange for the gift of *E. coli* MG1655, to Aman Soni for writing binned averaging code and to Sunish Radhakrishnan for access to phase-contrast microscopy and discussions on bacterial growth and microscopy.

## Author contributions

Tanvi Kale wrote the code, performed the experiments and analyzed the data. Dhruv Khatri wrote the analysis code and optimized the analysis. Chaitanya Athale supervised the project, wrote the paper and obtained funding support.

## Supplemetary material

**Video SV1:**
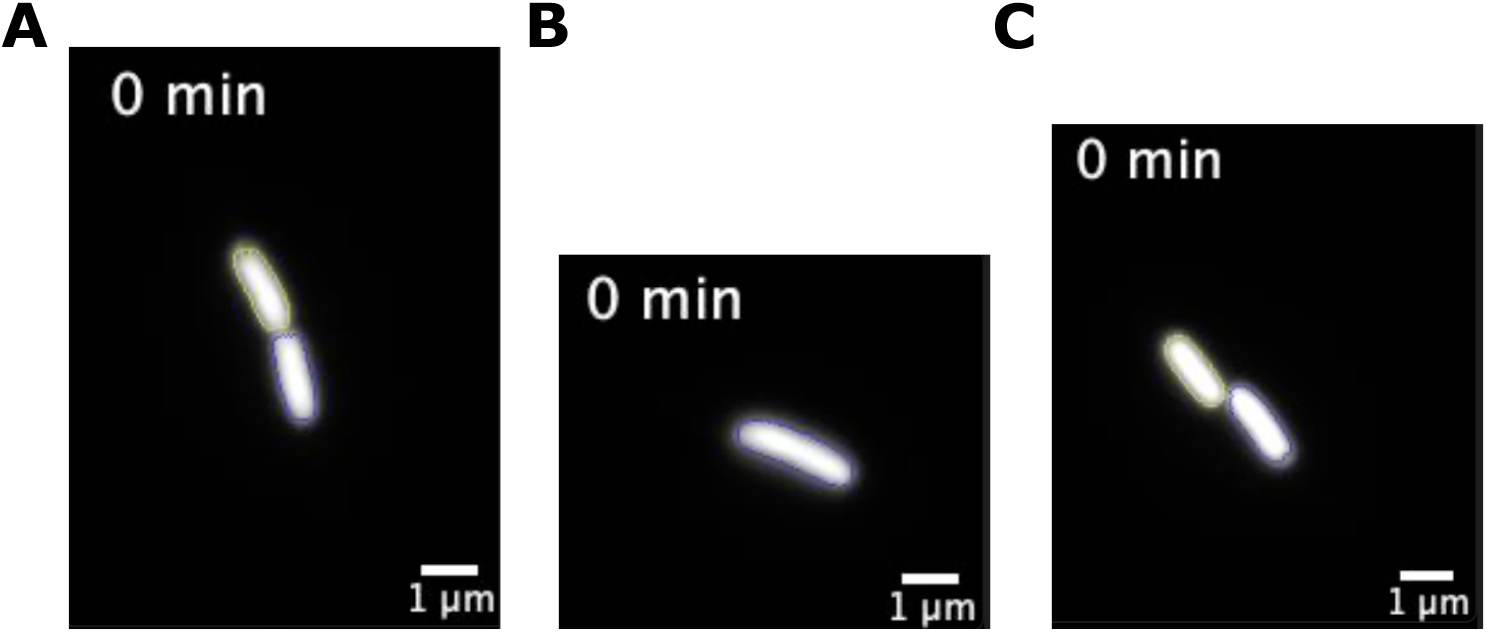
Single cell growth dynamics based on GFP labelled *E. coli* on nutrient agarose pads. The three representative movies (left to right) are of *E. coli* MG1655 cells expressing GFP grown on LB agarose pads at 37°C. Individual cells were detected (colored outlines) using an in-house developed image analysis pipeline as described in the Materials and Methods section. Time between frames: 1 minute, total time: 60 minutes. Time: minutes, scale-bar: 1 *μ*m.

